# Extremely potent human monoclonal antibodies from convalescent Covid-19 patients

**DOI:** 10.1101/2020.10.07.328302

**Authors:** Emanuele Andreano, Emanuele Nicastri, Ida Paciello, Piero Pileri, Noemi Manganaro, Giulia Piccini, Alessandro Manenti, Elisa Pantano, Anna Kabanova, Marco Troisi, Fabiola Vacca, Dario Cardamone, Concetta De Santi, Linda Benincasa, Chiara Agrati, Maria Rosaria Capobianchi, Concetta Castilletti, Arianna Emiliozzi, Massimiliano Fabbiani, Francesca Montagnani, Lorenzo Depau, Jlenia Brunetti, Luisa Bracci, Emanuele Montomoli, Claudia Sala, Giuseppe Ippolito, Rino Rappuoli

**Affiliations:** Monoclonal Antibody Discovery (MAD) Lab, Fondazione Toscana Life Sciences, Siena, Italy; VisMederi S.r.l, Siena, Italy; VisMederi Research S.r.l., Siena, Italy; National Institute for Infectious Diseases Lazzaro Spallanzani, IRCCS, Rome, Italy; Department of Medical Biotechnologies, University of Siena, Siena, Italy; Department of Medical Sciences, Infectious and Tropical Diseases Unit, University Hospital of Siena, Siena, Italy; Department of Molecular and Developmental Medicine, University of Siena, Siena, Italy; Faculty of Medicine, Imperial College, London, United Kingdom; Department of Biotechnology, Chemistry and Pharmacy, University of Siena, Siena, Italy; University of Turin, Turin, Italy; Tumour Immunology Unit, Fondazione Toscana Life Sciences, Siena, Italy; MedBiotech Hub and Competence Center, Department of Medical Biotechnologies, University of Siena

## Abstract

Human monoclonal antibodies are safe, preventive and therapeutic tools, that can be rapidly developed to help restore the massive health and economic disruption caused by the Covid-19 pandemic. By single cell sorting 4277 SARS-CoV-2 spike protein specific memory B cells from 14 Covid-19 survivors, 453 neutralizing antibodies were identified and 220 of them were expressed as IgG. Up to 65,9% of monoclonals neutralized the wild type virus at a concentration of >500 ng/mL, 23,6% neutralized the virus in the range of 100 - 500 ng/mL and 9,1% had a neutralization potency in the range of 10 - 100 ng/mL. Only 1,4% neutralized the authentic virus with a potency of 1-10 ng/mL. We found that the most potent neutralizing antibodies are extremely rare and recognize the RBD, followed in potency by antibodies that recognize the S1 domain, the S-protein trimeric structure and the S2 subunit. The three most potent monoclonal antibodies identified were able to neutralize the wild type and D614G mutant viruses with less than 10 ng/mL and are good candidates for the development of prophylactic and therapeutic tools against SARS-CoV-2.

**One Sentence Summary:** Extremely potent neutralizing human monoclonal antibodies isolated from Covid-19 convalescent patients for prophylactic and therapeutic interventions.

## INTRODUCTION

The impact of the SARS-CoV-2 pandemic, with more than 35 million cases, over 1 million deaths, 5 trillion impact on the gross domestic product (GDP) and 45 million people filing unemployment in the United States alone, is unprecedented (Aratani, 2020).

In the absence of drugs or vaccines, non-pharmaceutical interventions such as social distancing and quarantine have been the only way to contain the spread of the virus. These interventions showed to be efficient when properly implemented but not all countries were able to do so showing the limits of these strategies. The urgency to develop vaccines and therapies is extremely high. The effort to develop vaccines is unprecedented and fortunately in October 2020 we already have several vaccines in advanced Phase III efficacy trials and many others in earlier phase of development. In spite of the big effort, it is predictable this wave of infection will continue to spread globally and it is likely to be followed by additional waves in the next few years until herd immunity, acquired by vaccination or by natural infection, will constrain the circulation of the virus. It is therefore imperative to develop in parallel both vaccines and therapeutic tools to face the next waves of SARS-CoV-2 infections as soon as possible. Among the many therapeutic options available, human monoclonal antibodies (mAbs) are the ones that can be developed in the shortest period of time. In fact, the extensive clinical experience with the safety of more than 50 commercial mAbs approved to treat cancer, inflammatory and autoimmune, disorders provides high confidence on their safety. These advantages, combined with the urgency of the SARS-CoV-2 pandemic, support and justify an accelerated regulatory pathway. In addition, the long industrial experience in developing and manufacturing mAbs decreases the risks usually associated with the technical development of investigational products. Finally, the incredible technical progress in the field allows to shorten the conventional timelines and go from discovery to proof of concept trials in 5-6 months (Kelley, 2020). Indeed, in the case of Ebola, mAbs were the first therapeutic intervention recommended by the World Health Organization (WHO) and they were developed faster than vaccines or other drugs (Kupferschmidt, 2019).

The SARS-CoV-2 spike glycoprotein (S-protein) has a pivotal role in viral pathogenesis and it is considered the main target to elicit potent neutralizing antibodies and the focus for the development of therapeutic and prophylactic tools against this virus (Walls et al., 2020, Tay et al., 2020). Indeed, SARS-CoV-2 entry into host cells is mediated by the interaction between S-protein and the human angiotensin converting enzyme 2 (ACE2) (Wang et al., 2020b, Walls et al., 2020). The S-protein is a trimeric class I viral fusion protein which exists in a metastable prefusion conformation and in a stable postfusion state. Each S-protein monomer is composed of two distinct regions, the S1 and S2 subunits. Structural rearrangement occurs when the receptor binding domain (RBD) present in the S1 subunit binds to the host cell membrane. This interaction destabilizes the prefusion state of the S-protein triggering the transition into the postfusion conformation which in turn results in the ingress of the virus particle into the host cell (Wrapp et al., 2020). Single-cell RNA-sequencing analysis revealed that ACE2 expression was ubiquitous in different human organs have suggesting that SARS-CoV-2, through the S-protein, can invade human cells in different major physiological systems including the respiratory, cardiovascular, digestive and urinary systems, thus enhancing the possibility of spreading and infection (Zou et al., 2020).

During the first few months of this pandemic, several groups have been active in isolating and characterizing human monoclonal antibodies from Covid-19 convalescent patients or from humanized mice and some of them have been progressing quickly to clinical trials for the prevention and cure of SARS-Cov-2 infection (Shi et al., 2020, Hansen et al., 2020, Wang et al., 2020a, Pinto et al., 2020, Zost et al., 2020, Rogers et al., 2020, Andreano et al., 2020). So far, most of the work on human monoclonal antibodies against SARS-CoV-2 started from one or few patients and it allowed to successfully isolate and characterize the first interesting antibodies with moderate neutralization potency. In many cases an hundreds of nanograms to micrograms of antibodies were required to neutralize the virus *in vitro* therefore grams of antibodies will be needed per single patient. In this scenario intravenous delivery results to be the only possible administration route for their therapeutic and prophylactic use. Only recently very potent human antibodies have been isolated and these can be considered for intramuscular or subcutaneous administration. A striking example is a monoclonal antibody against RSV which delivered intramuscularly to premature babies has shown very promising results in clinical settings (Griffin et al., 2020). Furthermore, giving the initial rush in isolating potential antibody candidates for clinical development, the whole picture on different types of neutralizing antibodies generated after infection is not yet clear.

To identify and characterize potent mAbs against SARS-CoV-2, we isolated over 4,200 S-protein specific-memory B cells (MBCs) derived from 14 Covid-19 convalescent. From the screening of thousands of B cells, three extremely potent monoclonal antibodies were identified and are excellent candidates for further development.

## RESULTS

### Isolation and characterization of S-protein specific antibodies from SARS-CoV-2 convalescent patients

To retrieve mAbs specific for SARS-CoV-2 S-protein, peripheral blood mononuclear cells (PBMCs) from fourteen convalescent patients enrolled in this study were collected and stained with fluorescent labelled S-protein trimer to identify antigen specific memory B cells (MBCs). Fig. 1 summarizes the overall experimental strategy. The gating strategy described in Fig. S1 was used to single cell sort into 384-well plates IgG^+^ and IgA^+^ MBCs binding to the SARS-CoV-2 S-protein trimer in its prefusion conformation. The sorting strategy aimed to specifically identify class-switched MBCs (CD19^+^CD27^+^IgD^-^IgM^-^) to identify only memory B lymphocytes that went through maturation processes. A total of 4,277 S-protein-binding MBCs were successfully retrieved with frequencies ranging from 0,17% to 1,41% (Table 1). Following the sorting procedure, S-protein^+^ MBCs were incubated over a layer of 3T3-CD40L feeder cells in the presence of IL-2 and IL-21 stimuli for two weeks to allow natural production of immunoglobulins (10). Subsequently, MBC supernatants containing IgG or IgA were tested for their ability to bind either the SARS-CoV-2 S-protein trimer in its prefusion conformation or the S-protein S1 + S2 subunits (Fig. 2A - B) by enzyme linked immunosorbent assay (ELISA). A panel of 1,731 mAbs specific for the SARS-CoV-2 S-protein were identified showing a broad range of signal intensities (Table 1).

**Fig. 1.**
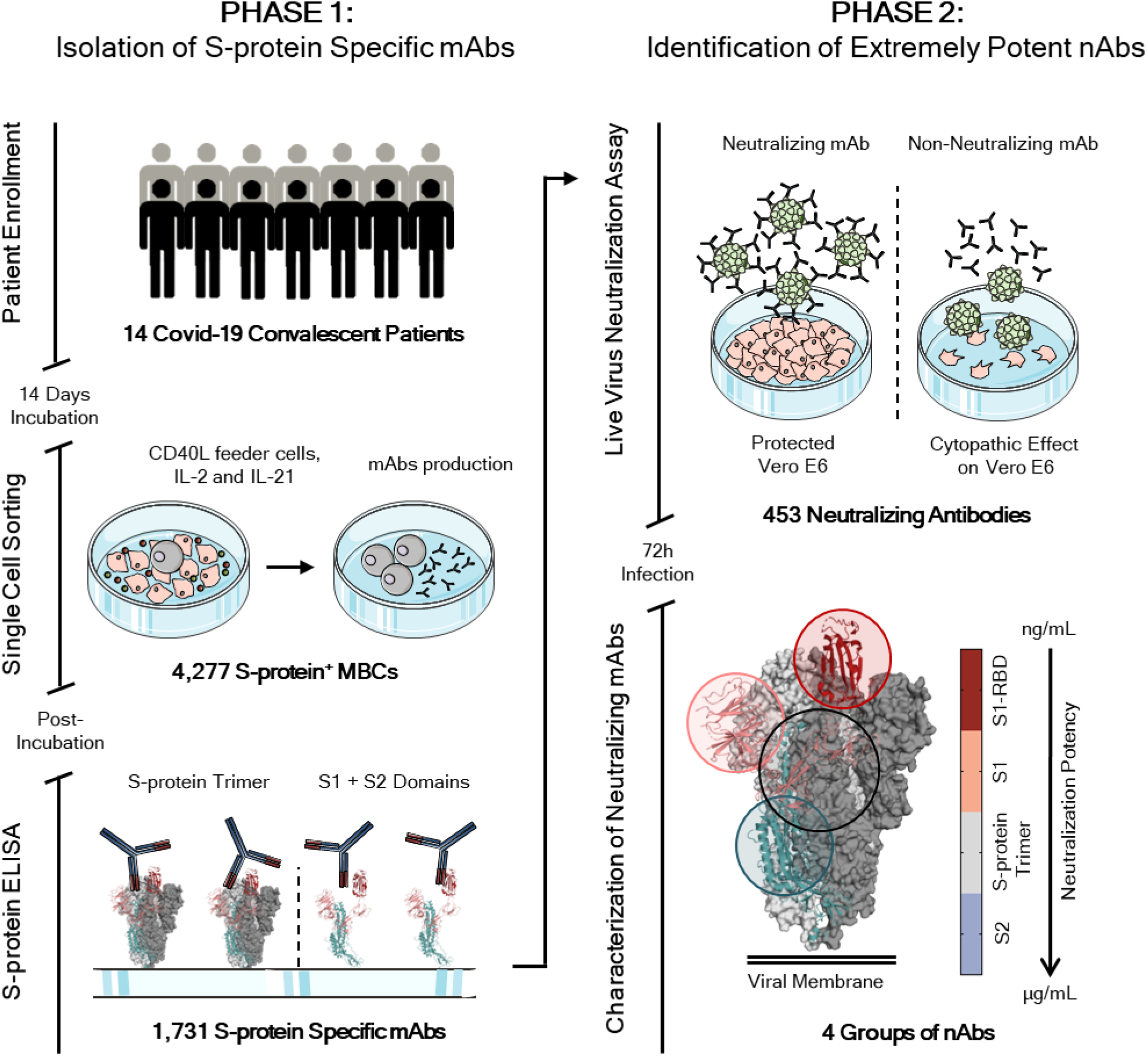
Workflow and timeline for SARS-CoV-2 neutralizing antibodies identification. The overall scheme shows two different phases for neutralizing antibodies (nAbs) identification. The phase 1 consisted in the enrolment of Covid-19 patients (14) from which PBMC was isolated. Memory B cells were single cell sorted (N= 4,277) and after 2 weeks of incubation the antibodies screened for their binding specificity against the S-protein trimer and S1/S2 domains. Once S-protein specific monoclonal antibodies were identified (N=1,731) the phase 2 started. All specific mAbs were tested *in vitro* to evaluate their neutralization activity against the live virus and 453 nAbs were identified. nAbs showing different binding profiles on the S-protein surface were selected for further functional characterization.

**Table 1.** Covid-19 convalescent patients summary. The table summarizes all the information collected per each donor such as percentage of S-protein specific memory B cells (MBCs), total number of S-protein specific MBCs sorted, S-protein specific mAbs, percentage of neutralizing mAbs and total number of nAbs identified.

| Subject ID | Antigen Specific MBCs (%) | S-protein<sup>+</sup> MBCs Sorted | S-protein<sup>+</sup> mAbs (ELISA) | Neutralization of binding (%) | SARS-CoV-2 Neutralizing mAbs |
| --- | --- | --- | --- | --- | --- |
| PT-004 | 1,41 | 367 | 158 | 61 | 13 |
| PT-005 | 0,55 | 108 | 33 | 3 | 4 |
| PT-006 | 0,32 | 99 | 34 | 3 | 4 |
| PT-008 | 0,46 | 374 | 148 | 24 | 48 |
| PT-009 | 0,81 | 136 | 61 | 8 | 12 |
| PT-010 | 0,34 | 97 | 39 | 4 | 3 |
| PT-012 | 0,86 | 359 | 132 | 15 | 14 |
| PT-014 | 0,52 | 308 | 165 | 29 | 53 |
| PT-041 | 0,70 | 316 | 165 | 47 | 77 |
| PT-100 | 0,76 | 506 | 230 | 55 | 59 |
| PT-101 | 0,55 | 682 | 219 | 26 | 66 |
| PT-102 | 0,99 | 247 | 145 | 29 | 42 |
| PT-103 | 0,34 | 286 | 129 | 25 | 54 |
| PT-188 | 0,17 | 299 | 72 | 8 | 4 |
| Total (N) |  | 4,277 | 1,731 | 339 | 453 |
| Total (%) |  |  | 40,4 | 19,6 | 26,2 |

**Fig. 2.**
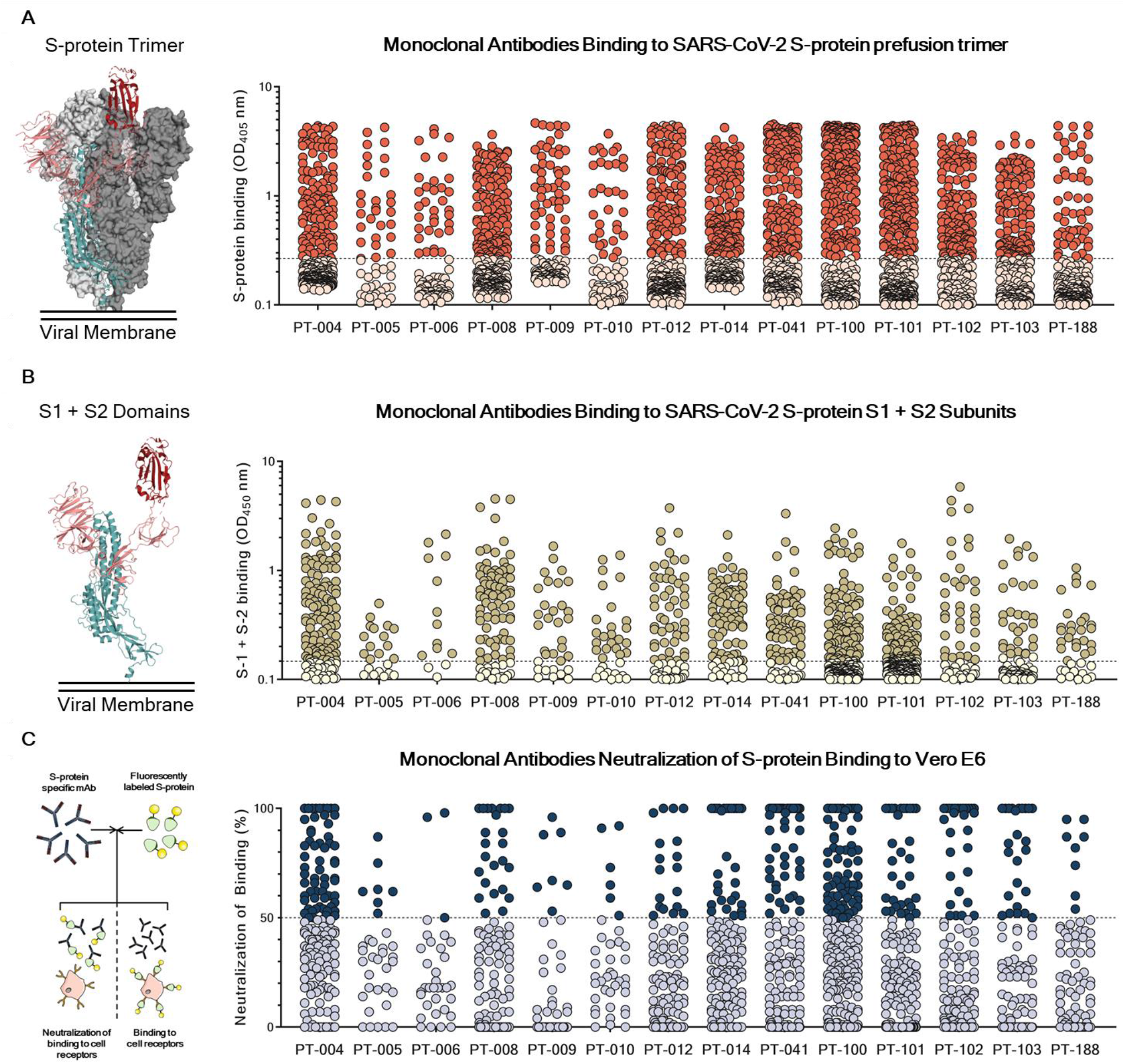
Identification and characterization of SARS-CoV-2 S-protein specific mAbs. (A) The graph shows supernatants tested for binding to the SARS-CoV-2 S-protein stabilized in its prefusion conformation. Threshold of positivity has been set as two times the value of the blank (dotted line). Red dots represent mAbs which bind to the S-protein while pink dots represent mAbs which do not bind. (B) The graph shows supernatants tested for binding to the SARS-CoV-2 S-protein S1 + S2 subunits. Threshold of positivity has been set as two times the value of the blank (dotted line) Darker dots represent mAbs which bind to the S1 + S2 while light yellow dots represent mAbs which do not bind. (C) The graph shows supernatants tested by NoB assay. Threshold of positivity has been set as 50% of binding neutralization (dotted line). Dark blue dots represent mAbs able to neutralize the binding between SARS-CoV-2 and receptors on Vero E6 cells, while light blue dots represent non-neutralizing mAbs. The total number (N) of S-protein specific supernatants screened by NoB assay is shown for each donor.

### Identification of S-protein specific mAbs able to neutralize SARS-CoV-2

The 1,731 supernatants containing S-protein specific mAbs, were screened *in vitro* for their ability to block the binding of the S-protein to Vero E6 cell receptors. and for their ability to neutralize authentic SARS-CoV-2 virus by *in vitro* microneutralization assay. In the neutralization of binding (NoB) assay, 339 of the 1,731 tested (19,6%) S-protein specific mAbs were able to neutralize the antigen/receptor binding showing a broad array of neutralization potency ranging from 50% to over 100% (Table 1 and Fig. 2C).

As for the authentic virus neutralization assay, supernatants containing naturally produced IgG or IgA were tested for their ability to protect the layer of Vero E6 cells from the cytopathic effect triggered by SARS-CoV-2 infection (Fig. S2). To increase the throughput of our approach, supernatants were tested at a single point dilution and to increase the sensibility of our first screening a viral titer of 25TCID_50_ was used. For this first screening mAbs were classified as neutralizing, partially neutralizing and not-neutralizing mAbs based on their inability to protect Vero E6 cells from infection, or to their ability to partially or completely prevent the cytopathic effect. Out of 1,731 mAbs tested in this study, a panel of 453 (26,2%) mAbs neutralized the live virus and prevented infection of Vero E6 cells (Table 1). The percentage of partially neutralizing mAbs and neutralizing mAbs (nAbs) identified in each donor was extremely variable ranging from 2,6 - 29,7% and 2,8 - 26,4% respectively (Fig. 3A and Table S1). The majority of nAbs were able to specifically recognize the S-protein S1 domain (57,5%; N=244) while 7,3% (N=53) of nAbs were specific for the S2 domain and 35,2% (N=156) did not recognize single domains but only the S-protein in its trimeric conformation (Fig. 3B; Table S2). From the panel of 453 nAbs, we recovered the heavy and light chain amplicons of 220 nAbs which were expressed as full length IgG1 using the transcriptionally active PCR (TAP) approach to characterize their neutralization potency. The vast majority of nAbs identified (65,9%; N=145) had a low neutralizing potency and required more than 500 ng/ml to achieve an IC100. A smaller fraction of the antibodies had an intermediate neutralizing potency (23,6%; N=52) requiring between 100 and 500 ng/ml to achieve the IC_100_, while 9,1% (N=20) required between 10 and 100 ng/ml. Finally, only 1,4% (N=3) of the expressed nAbs were classified as extremely potently nAbs, showing an IC100 lower than 10 ng/mL (Fig. 3C - D; Table S3).

**Fig. 3.**
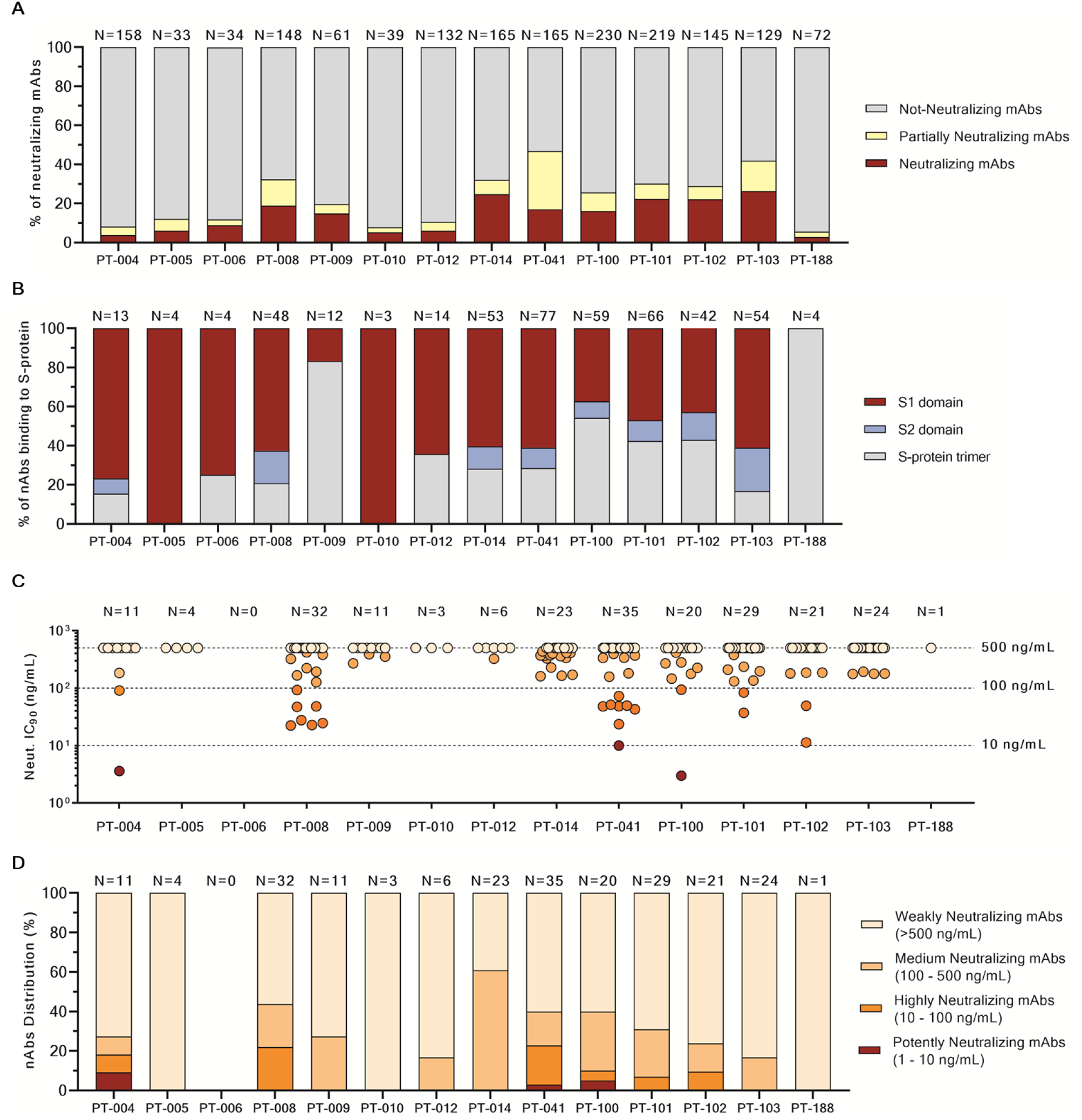
Identification of SARS-CoV-2 S-protein specific neutralizing antibodies. (A) The bar graph shows the percentage of not-neutralizing (gray), partially neutralizing (pale yellow) and neutralizing mAbs (dark red) identified per each donor. The total number (N) of antibodies tested per individual is shown on top of each bar. (B) The bar graph shows the distribution of nAbs binding to different S-protein domains. In dark red, light blue and gray are shown antibodies binding to the S1-domain, S2 domain and S-protein trimer respectively. The total number (N) of antibodies tested per individual is shown on top of each bar. (C) The graph shows the neutralization potency of each nAb tested once expressed as recombinant full-length IgG1. Dashed lines show different ranges of neutralization potency (500 – 100 – 10 ng/mL). Dots were colored based on their neutralization potency and were classified as weakly neutralizing (>500 ng/mL; pale orange), medium neutralizing (100 – 500 ng/mL; orange), highly neutralizing (10 – 100 ng/mL; dark orange) and extremely neutralizing (1 – 10 ng/mL; dark red). The total number (N) of antibodies tested per individual is shown on top of each graph. (D) The bar graph shows the percentage of nAbs with different neutralization potencies. nAbs were classified as weakly neutralizing (>500 ng/mL; pale orange), medium neutralizing (100 – 500 ng/mL; orange), highly neutralizing (10 – 100 ng/mL; dark orange) and extremely neutralizing (1 – 10 ng/mL; dark red). The total number (N) of antibodies tested per individual is shown on top of each bar.

### Classification of SARS-CoV-2 nAbs

Based on the first round of screening, 14 nAbs were selected for further characterization. All nAbs were able to bind the SARS-CoV-2 S-protein in its trimeric conformation (Fig. 4A). The mAbs named J08, I14, F05, G12, C14, B07, I21, J13 and D14 were also able to specifically bind the S1 domain (Fig. 4B). The nAbs named H20, I15, F10 and F20 were not able to bind single S1 or S2 domains but only the S-protein in its trimeric state, while the nAb L19 bound only the S2 subunit (Fig. 4B - C). Among the group of S1 specific nAbs only J08, I14, F05, G12, C14 and B07 were able to bind the S1 receptor binding domain (RBD) and to strongly inhibit the interaction between the S-protein and Vero E6 receptors (Fig. 2C; Fig. S3; Table 2). On the other hand I21, J13 and D14, despite showing S1 binding specificity, did not show any binding to the RBD and NoB activity (Fig S3 and Table 2). Based on this description four different groups of nAbs against SARS-CoV-2 were identified. The first group (Group I) is composed by S1-RBD specific nAbs (J08, I14, F05, G12 and C14) and showed extremely high neutralization potency against both the WT and D614G live viruses ranging from 3,91 to 157,49 ng/mL (Fig. 4D - F; Table 2). The second group (Group II) is composed by S1 nAbs that did not bind the RBD (B07, I21, J13 and D14). These antibodies also showed good neutralization potency ranging from 49,61 to 500 ng/mL (Fig. 4D - F; Table 2) but inferior to S1-RBD directed nAbs. Antibodies belonging to Group I and II showed picomolar affinity to the S-protein with a KD ranging from 0.78 to 6.30 E^−10^M (Fig. S4). The third group (Group III) is composed by antibodies able to bind the S-protein only in its whole trimeric conformation (H20, I15, F10 and F20). Antibodies belonging to this group showed lower affinity to the S-protein (KD 7.57 E^−8^M - 6.40 E^−9^M) compared to Group I and II nAbs and medium neutralization potencies ranging from 155,02 to 492,16 ng/mL (Fig. 4D - F; Table 2; Fig. S4). The fourth and final group (Group IV) composed only by L19 nAb and recognized exclusively the S2 domain of the S-protein showing the lowest neutralization potency with 19,8 μg/mL and 12,5 μg/mL for the authentic WT and D614G strain respectively (Fig. 4D-F; Table 2).

**Fig. 4.**
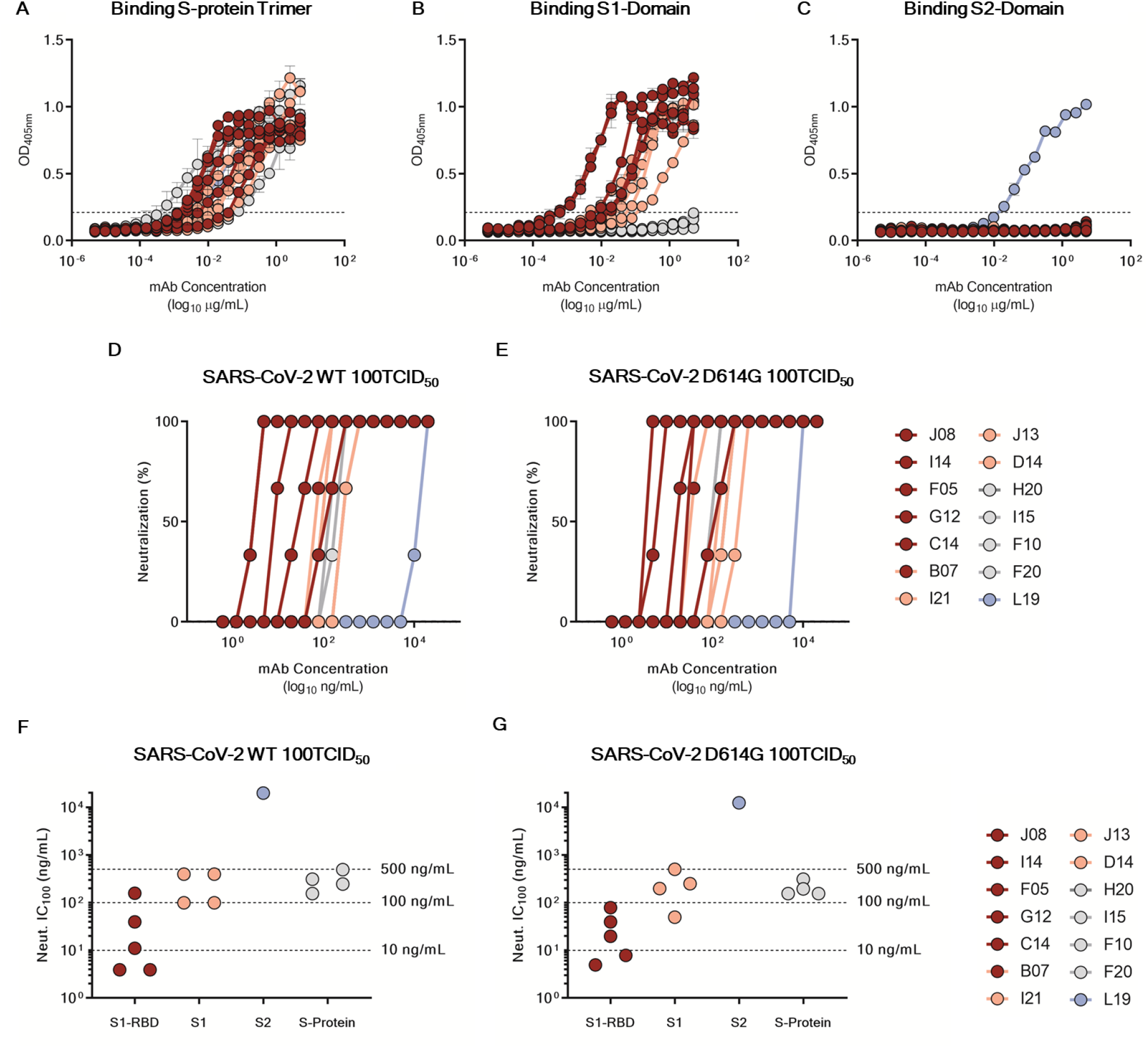
Functional characterization of potent SARS-CoV-2 S-protein specific nAbs. (A - B - C) Graphs show binding curves to the S-protein in its trimeric conformation, S1-domain and S2-domain. Mean ± SD of technical triplicates are shown. Dashed lines represent the threshold of positivity (D – E) Neutralization curves for selected antibodies were shown as percentage of viral neutralization against the authentic SARS-CoV-2 wild type and D614G strains. Data are representative of technical triplicates. (F – G) Neutralization potency of fourteen selected antibodies against the authentic SARS-CoV-2 wild type and D614G strains. Dashed lines show different ranges of neutralization potency (500 – 100 – 10 ng/mL). In all graphs selected antibodies are shown in dark red, pink, gray and light blue based on their ability to recognize the SARS-CoV-2 S1-RBD, S1-domain, S-protein trimer only and S2-domain respectively.

**Table 2.** Characteristics of selected neutralizing antibodies. The table summarizes the binding specificity, affinity, NoB and neutralization features for the fourteen neutralizing antibodies selected in this study.

| mAb ID | Binding specificity | S-protein Affinity KD (M) | Neutralization of Binding (%) | Neutralization Titer IC100 (ng/mL)_WT | Neutralization Titer IC100 (ng/mL)_D614G |
| --- | --- | --- | --- | --- | --- |
| J08 | S-1 | 0,78E-10 | 100 | 3,91 | 7,81 |
| I14 | S-1 | 6,30E-10 | 99 | 11,05 | 19,69 |
| F05 | S-1 | 3,04E-10 | 89 | 3,91 | 4,92 |
| G12 | S-1 | 4,36E-10 | 100 | 39,37 | 39,37 |
| C14 | S-1 | 1,19E-10 | 100 | 157,49 | 78,75 |
| B07 | S-1 | 4,60E-10 | 100 | 99,21 | 49,61 |
| I21 | S-1 | N/A | 0 | 99,21 | 198,43 |
| J13 | S-1 | N/A | 0 | 396,85 | 500 |
| D14 | S-1 | N/A | 0 | 396,85 | 250 |
| H20 | S-protein | N/A | 0 | 492,16 | 310,04 |
| I15 | S-protein | 6,40E-09 | 46 | 310,04 | 155,02 |
| F10 | S-protein | 3,03E-08 | 45 | 155,02 | 195,31 |
| F20 | S-protein | 7,57E-08 | 21 | 246,08 | 155,02 |
| L19 | S-2 | N/A | 15 | 19842,51 | 12500 |

## DISCUSSION

This work describes a systematic screening of memory B cells from convalescent people to identify extremely potent human monoclonal antibodies against the spike protein of the SARS-CoV-2 virus, to be used for prevention and therapy of Covid-19. We found that approximately 10% of the total B cells against the spike protein produce neutralizing antibodies and these can be divided into 4 different groups recognizing the S1-RBD, S1-domain, S2-domain and the S-protein trimer. We found that the most potent neutralizing antibodies are extremely rare and that they recognize the RBD, followed in potency by the antibodies recognizing the S1 domain, the trimeric structure and the S2 subunit. The L19 antibody against the S2 subunit, which had the lowest neutralizing potency, is representative of several other S2 specific antibodies identified in the preliminary screening. From these data we conclude that in convalescent patients most of the observed neutralization titers are mediated by the antibodies with medium-high neutralizing potency. Indeed, the extremely potent antibodies and the antibodies against the S2 subunit are unlikely to contribute to the overall neutralizing titers because they are too rare and too poor neutralizers respectively to be able to make the difference. The observed antibody repertoire of convalescent patients may be a consequence of the loss of Bcl-6-expressing follicular helper T cells and the loss of germinal centers in Covid-19 patients which may limit and constrain the B cell affinity maturation (Kaneko et al., 2020). It is therefore important to perform similar studies following vaccination as it is likely that the repertoire of neutralizing antibodies induced by vaccination may be different from the one described here.

Out of 453 neutralizing antibodies that were tested and characterized three showed extremely high neutralization potency against both the initial SARS-CoV-2 strain isolated in Wuhan and the D614G variant currently spread worldwide. During the last few months several groups reported the isolation, structure and passive protection in animal models of neutralizing antibodies against SARS-CoV-2. Most of these studies, with few exceptions report antibodies which require from 20 to several hundreds more ng/mL to neutralize 50% of the virus *in vitro*. These antibodies are potentially good for therapy. However, they will require a high dosage which will result in elevated cost of goods, low capacity to numbers large quantities of doses and intravenous infusion.

The extremely potent candidates described in our study will allow to use small quantities of antibodies to reach the prophylactic and therapeutic dosage and as consequence decrease the cost of goods and implement sustainable development and manufacturability. This solution may increase the number of doses produced annually and therefore increase antibodies availability in high income countries as well as low-and middle-income countries (LMICs). Our work combined with institutions such as the Coalition for Epidemic Preparedness Innovations (CEPI) can be used synergically to bring SARS-CoV-2 prophylactic and therapeutic tools worldwide and help the fight against Covid-19 pandemic.

## ACKNOWLEDGMENTS

We wish to thank Fondazione Toscana Life Sciences in the persons of Dr. Fabrizio Landi and Dr. Andrea Paolini and the whole Administration for their incredible help and support. In particular we would like to thank Mr. Francesco Senatore, Mrs. Laura Canavacci and Mrs. Cinzia Giordano for their support in preparing all the documents needed for the ethical approval of the clinical studies carried out within this project.

This work was possible thanks to the technology established in our European Research Council (ERC) funded laboratory through the ERC Advanced grant vAMRes which allowed us to isolate mAbs from vaccinated and/or convalescent patients to tackle the global threat posed by antimicrobial resistance.

We wish to thank the National Institute for Infectious Diseases, IRCCS, Lazzaro Spallanzani Rome (IT) and the Azienda Ospedaliera Universitaria Senese, Siena (IT), for providing blood samples from Covid-19 convalescent donors under studies approved by local ethic committees. We also wish to thank all the nursing staff who chose to cooperate for blood withdrawal and all the donors who decided to participate in this study.

We would like to thank the whole GSK Vaccines Pre-clinical Evidence Generation and Assay - Immunolgy function led by Dr. Oretta Finco for their availability and support as well as Mrs. Simona Tavarini, Mrs. Chiara Sammicheli, Dr. Monia Bardelli, Dr. Michela Brazzoli, Dr. Elisabetta Frigimelica, Dr. Erica Borgogni and Dr. Elisa Faenzi for sharing their expertise, extreme availability and technical support. We would also like to thank Dr. Mariagrazia Pizza and Dr. Simone Pecetta for initial insightful advice and discussions on this project.

We would like to thank Dr. Jason McLellan and his team for generously providing the SARS-CoV-2 S-protein stabilized in its prefusion conformation used in this study. Furthermore, we would like to thank Dr. Daniel Wrapp and Dr. Nianshuang Wang for the precious information and suggestions.

We gratefully acknowledge the Collaborators Members of INMI COVID-19 study group and all the other members of the Covid-19 UNIT and of the Covid-19 TEAM, AOUS (Azienda Ospedaliera Universitaria Senese, Siena, Italy).

This publication was supported by funds from the “Centro Regionale Medicina di Precisione” and by all the people who answered the call to fight with us the battle against SARS-CoV-2 with their kind donations on the platform ForFunding (https://www.forfunding.intesasanpaolo.com/DonationPlatform-ISP/nav/progetto/id/3380).

This publication was supported by the European Virus Archive goes Global (EVAg) project, which has received funding from the European Union’s Horizon 2020 research and innovation programme under grant agreement No 653316.

This publication was supported by the COVID-2020-12371817 project, which has received funding from the Italian Ministry of Health.

## AUTHOR CONTRIBUTIONS

Emanuele Andreano, Ida Paciello isolated single memory B cells and identified S-protein specific mAbs (cell sorting and ELISA).

Piero Pileri, Noemi Manganaro, Elisa Pantano performed NoB assay and produced recombinant spike protein.

Emanuele Andreano, Giulia Piccini, Alessandro Manenti and Emanuele Montomoli performed ELISA and viral neutralization assay.

Marco Troisi, Fabiola Vacca, recovered heavy and light chain amplicons and expressed recombinant antibodies.

Ida Paciello and Piero Pileri performed RBD-antibodies binding assay.

Concetta De Santi, Dario Cardamone, Anna Kabanova contributed to the characterization of positive memory B cells.

Emanuele Nicastri, Chiara Agrati, Concetta Castilletti, Francesca Montagnani, Arianna Emiliozzi, Massimiliano Fabbiani, Maria Rosaria Capobianchi enrolled patients and isolated PBMCs.

Luisa Bracci, Jlenia Brunetti and Lorenzo Depau performed surface plasmon resonance assay to identify antibody affinities.

Claudia Sala, Giuseppe Ippolito, Rino Rappuoli coordinated the project.

## METHOD DETAILS

### Enrollment of SARS-COV-2 convalescent donors and human sample collection

This work results from a collaboration with the National Institute for Infectious Diseases, IRCCS - Lazzaro Spallanzani Rome (IT) and Azienda Ospedaliera Universitaria Senese, Siena (IT) that provided samples from SARS-CoV-2 convalescent donors who gave their written consent. The study was approved by local ethics committees (Parere 18_2020 in Rome and Parere 17065 in Siena) and conducted according to good clinical practice in accordance with the declaration of Helsinki (European Council 2001, US Code of Federal Regulations, ICH 1997). This study was unblinded and not randomized.

### Human peripheral blood mononuclear cells (PBMCs) isolation from SARS-CoV-2 convalescent donors

Peripheral blood mononuclear cells (PBMCs) were isolated from heparin-treated whole blood by density gradient centrifugation (Lympholyte-H; Cedarlane). After separation, PBMC were: i) frozen in liquid nitrogen at concentration of 10 x 10^6^ PBMC/vial using 10% DMSO in heat-inactivated fetal bovine serum (FBS) or ii) resuspended in RPMI 1640 (EuroClone) supplemented with 10% FBS (EuroClone), 2 mmol/L L-glutamine, 2 mmol/L penicillin, and 50 μg/mL streptomycin (EuroClone). Cells were cultured for 18 hour at 37°C with 5% CO_2_. Blood samples were screened for SARS-CoV-2 RNA and for antibodies against HIV, HBV and HCV.

### Expression and purification of SARS-CoV-2 S-protein prefusion trimer

The expression vector coding for prefusion S ectodomain (kind gift of Dr. Jason Mc Lellan) was used to transiently transfect Expi293F cells (Thermo Fisher #A14527) using Expifectamine (Thermo Fisher # A14525). The protein was purified from filtered cell supernatants using NiNTA resin (GE Healtcare #11-0004-58), eluted with 250 mM Imidazole (Sigma Aldrich #56750), dialyzed against PBS, and then stored at 4°C prior to use.

### Single cell sorting of SARS-CoV-2 S-protein^+^ memory B cells

Human peripheral blood mononuclear cells (PBMCs) from SARS-CoV-2 convalescent donors were stained with Live/Dead Fixable Aqua (Invitrogen; Thermo Scientific) in 100 μL final volume diluted 1:500 at room temperature (RT). After 20 min incubation cells were washed with phosphate buffered saline (PBS) and unspecific bindings were saturated with 50 μL of 20% rabbit serum in PBS. Following 20 min incubation at 4°C cells were washed with PBS and stained with SARS-CoV-2 S-protein labeled with Strep-Tactin^®^XT DY-488 (iba-lifesciences cat# 2-1562-050) for 30 min at 4°C. After incubation the following staining mix was used CD19 V421 (BD cat# 562440), IgM PerCP-Cy5.5 (BD cat# 561285), CD27 PE (BD cat# 340425), IgD-A700 (BD cat# 561302), CD3 PE-Cy7 (BioLegend cat# 300420), CD14 PE-Cy7 (BioLegend cat# 301814), CD56 PE-Cy7 (BioLegend cat# 318318) and cells were incubated at 4°C for additional 30 min. Stained MBCs were single cell-sorted with a BD FACSAria III (BD Biosciences) into 384-well plates containing 3T3-CD40L feeder cells and were incubated with IL-2 and IL-21 for 14 days as previously described (Huang et al., 2013).

### ELISA assay with S1 and S2 subunits of SARS-CoV-2 S-protein

The presence of S1- and S2-binding antibodies in culture supernatants of monoclonal S-protein-specific memory B cells was assessed by means of an ELISA assay implemented with the use of a commercial kit (ELISA Starter Accessory Kit, Catalogue No. E101; Bethyl Laboratories). Briefly, 384-well flat-bottom microtiter plates (Nunc MaxiSorp 384-well plates; Sigma-Aldrich) were coated with 25 μl/well of antigen (1:1 mix of S1 and S2 subunits, 1 μg/ml each; The Native Antigen Company, Oxford, United Kingdom) diluted in coating buffer (0.05 M carbonate-bicarbonate solution, pH 9.6), and incubated overnight at 4°C. The plates were then washed three times with 100 μl/well washing buffer (50 mM Tris Buffered Saline (TBS) pH 8.0, 0.05% Tween-20) and saturated with 50 μl/well blocking buffer containing Bovine Serum Albumin (BSA) (50 mM TBS pH 8.0, 1% BSA, 0.05% Tween-20) for 1 hour (h) at 37°C. After further washing, samples diluted 1:5 in blocking buffer were added to the plate. Blocking buffer was used as a blank. After an incubation of 1 h at 37°C, plates were washed and incubated with 25 μl/well secondary antibody (horseradish peroxidase (HRP)-conjugated goat anti-human IgG-Fc Fragment polyclonal antibody, diluted 1:10,000 in blocking buffer, Catalogue No. A80-104P; (Bethyl Laboratories) for 1 h at 37°C. After three washes, 25 μl/well TMB One Component HRP Microwell Substrate (Bethyl Laboratories) was added and incubated 10-15 minutes at RT in the dark. Color development was terminated by addition of 25 μl/well 0.2 M H2SO4. Absorbance was measured at 450 nm in a Varioskan Lux microplate reader (Thermo Fisher Scientific). The threshold for sample positivity was set at twice the optical density (OD) of the blank.

### ELISA assay with SARS-CoV-2 S-protein prefusion trimer

ELISA assay was used to detect SARS-CoV-2 S-protein specific mAbs and to screen plasma from SARS-CoV-2 convalescent donors. 384-well plates (Nunc MaxiSorp 384 well plates; Sigma Aldrich) were coated with 3μg/mL of streptavidin diluted in PBS and incubated at RT overnight. Plates were then coated with SARS-CoV-2 S-protein at 3μg/mL and incubated for 1h at room temperature. 50 μL/well of saturation buffer (PBS/BSA 1%) was used to saturate unspecific binding and plates were incubated at 37°C for 1h without CO_2_. Supernatants were diluted 1:5 in PBS/BSA 1%/Tween20 0,05% in 25 μL/well final volume and incubated for 1h at 37°C without CO_2_. 25 μL/well of alkaline phosphatase-conjugated goat anti-human IgG (Sigma-Aldrich) and IgA (Jackson Immuno Research) were used as secondary antibodies. In addition, twelve two-fold serial dilutions of plasma from SARS-CoV-2 infected patients were analyzed in duplicate. Plasma samples were diluted in PBS/BSA 1%/Tween20 0,05% (25 μL/well final volume; Starting Dilution 1:80) and incubated for 1h at 37°C without CO_2_. Next, 25 μL/well of alkaline phosphatase-conjugated goat anti-human IgG (Sigma-Aldrich) was added for 1h at 37°C without CO_2_. Wells were washed three times between each step with PBS/BSA 1%/Tween20 0.05%. PNPP (p-nitrophenyl phosphate) (Thermo Fisher) was used as soluble substrate to detect SARS-CoV-2 S-protein specific monoclonal antibodies and the final reaction was measured by using the Varioskan Lux Reader (Thermo Fisher Scientific) at a wavelength of 405 nm. Samples were considered as positive if OD at 405 nm (OD405) was two times the blank.

### SARS-CoV-2 virus and cell infection

African green monkey kidney cell line Vero E6 cells (American Type Culture Collection [ATCC] #CRL-1586) were cultured in Dulbecco’s Modified Eagle’s Medium (DMEM) - High Glucose (Euroclone, Pero, Italy) supplemented with 2 mM L-Glutamine (Lonza, Milano, Italy), penicillin (100 U/mL) - streptomycin (100 μg/mL) mixture (Lonza, Milano, Italy) and 10% Foetal Bovine Serum (FBS) (Euroclone, Pero, Italy). Cells were maintained at 37°C, in a 5% CO_2_ humidified environment and passaged every 3-4 days.

Wild type SARS CoV-2 2019 (2019-nCoV strain 2019-nCov/Italy-INMI1) virus was purchased from the European Virus Archive goes Global (EVAg, Spallanzani Institute, Rome). For virus propagation, sub-confluent Vero E6 cell monolayers were prepared in T175 flasks (Sarstedt) containing supplemented D-MEM high glucose medium. For titration and neutralization tests of SARS-CoV-2, Vero E6 were seeded in 96-well plates (Sarstedt) at a density of 1,5×10^4^ cells/well the day before the assay.

### Neutralization of Binding (NoB) Assay

To study the binding of the Covid-19 Spike protein to cell-surface receptor(s) we developed an assay to assess recombinant Spike protein specific binding to target cells and neutralization thereof. To this aim the stabilized Spike protein was coupled to Streptavidin-PE (eBioscience # 12-4317-87, Thermo Fisher) for 1h at room temperature and then incubated with VERO E6 cells. Binding was assessed by flow cytometry. The stabilized Spike protein bound VERO E6 cells with high affinity (data not shown). To assess the content of neutralizing antibodies in sera or in B-cell culture supernatants, two microliters of SARS-CoV-2 Spike-Streptavidin-PE at 5-10 μg/ml in PBS-5%FCS were mixed with two microliters of various dilutions of sera or B-cell culture supernatants in U bottom 96-well plates. After incubation at 37°C for 1 hr, 30×10^3^ Vero E6 cells suspended in two microliters of PBS 5% FCS were added and incubated for additional 45 min at 4°C. Non-bound protein and antibodies were removed and cell-bound PE-fluorescence was analyzed with a FACScantoII flow cytometer (Becton Dickinson). Data were analyzed using the FlowJo data analysis software package (TreeStar, USA). The specific neutralization was calculated as follows: NOB (%) = 1 - (Sample MFI value - background MFI value) / (Negative Control MFI value - background MFI value).

### Single cell RT-PCR and Ig gene amplification

From the original 384-well sorting plate, 5 μL of cell lysate was used to perform RT-PCR. Total RNA from single cells was reverse transcribed in 25 μL of reaction volume composed by 1 μL of random hexamer primers (50 ng/μL), 1 μL of dNTP-Mix (10 mM), 2 μL 0.1 M DTT, 40U/μL RNAse OUT, MgCl2 (25 mM), 5x FS buffer and Superscript^®^ IV reverse transcriptase (Invitrogen). Final volume was reached by adding nuclease-free water (DEPC). Reverse transcription (RT) reaction was performed at 42°C/10’, 25°C/10’, 50°C/60’ and 94°/5’. Heavy (VH) and light (VL) chain amplicons were obtained via two rounds of PCR. All PCR reactions were performed in a nuclease-free water (DEPC) in a total volume of 25 μL/well. Briefly, 4 μL of cDNA were used for the first round of PCR (PCRI). PCRI-master mix contained 10 μM of VH and 10 μM VL primer-mix, 10mM dNTP mix, 0,125 μL of Kapa Long Range Polymerase (Sigma), 1,5 μL MgCl2 and 5 μL of 5x Kapa Long Range Buffer. PCRI reaction was performed at 95°/3’, 5 cycles at 95°C/30”, 57°C/30”, 72°C/30” and 30 cycles at 95°C/30”, 60°C/30”, 72°C/30” and a final extension of 72°/2’. All nested PCR reactions (PCRII) were performed using 3,5 μL of unpurified PCRI product using the same cycle conditions. PCRII products were then purified by Millipore MultiScreen^®^ PCRμ96 plate according to manufacture instructions. Samples were eluted with 30 μl nuclease-free water (DEPC) into 96-well plates and quantify by Qubit Fluorometric Quantitation assay (Invitrogen).

### Cloning of variable region genes and recombinant antibody expression

Vector digestions were carried out with the respective restriction enzymes AgeI, SalI and Xho as previously described (Tiller et al., 2008, Wardemann and Busse, 2019). Briefly, 75 ng of IgH, Igλ and Igκ purified PCRII products were ligated by using the Gibson Assembly NEB into 25 ng of respective human Igγ1, Igκ and Igλ expression vectors. The reaction was performed into 5 μL of total volume. Ligation product was 10-fold diluted in nuclease-free water (DEPC) and used as template for transcriptionally active PCR (TAP) reaction which allowed the direct use of linear DNA fragments for *in vitro* expression. The entire process consists of one PCR amplification step, using primers to attach functional promoter (human CMV) and terminator sequences (SV40) onto the fragment PCRII products. TAP reaction was performed in a total volume of 25 μL using 5 μL of Q5 polymerase (NEB), 5 μL of GC Enhancer (NEB), 5 μL of 5X buffer,10 mM dNTPs, 0,125 μL of forward/reverse primers and 3 μL of ligation product. TAP reaction was performed by using the following cycles: 98°/2’, 35 cycles 98°/10”, 61°/20”, 72°/1’ and 72°/5’ as final extention step. TAP products were purified under the same PCRII conditions, quantified by Qubit Fluorometric Quantitation assay (Invitrogen) and used for transient transfection in Expi293F cell line using manufacturing instructions.

### Viral propagation and titration

The SARS-CoV-2 virus was propagated in Vero E6 cells cultured in DMEM high Glucose supplemented with 2% FBS, 100 U/mL penicillin, 100 μg/mL streptomycin. Cells were seeded at a density of 1×10^6^ cells/mL in T175 flasks and incubated at 37°C, 5% CO_2_ for 18-20 hours. The sub-confluent cell monolayer was then washed twice with sterile Dulbecco’s phosphate buffered saline (DPBS). Cells were inoculated with 3,5 mL of the virus properly diluted in DMEM 2% FBS at a multiplicity of infection (MOI) of 0.001, and incubated for 1h at 37°C in a humidified environment with 5% CO_2_. At the end of the incubation, 50 mL of DMEM 2% FBS were added to the flasks. The infected cultures were incubated at 37°C, 5% CO_2_ and monitored daily until approximately 80-90% of the cells exhibited cytopathic effect (CPE). Culture supernatants were then collected, centrifuged at 4°C at 1,600 rpm for 8 minutes to allow removal of cell debris, aliquoted and stored at −80°C as the harvested viral stock. Viral titers were determined in confluent monolayers of Vero E6 cells seeded in 96-well plates using a 50% tissue culture infectious dose assay (TCID_50_). Cells were infected with serial 1:10 dilutions (from 10-1 to 10-11) of the virus and incubated at 37°C, in a humidified atmosphere with 5% CO_2_. Plates were monitored daily for the presence of SARS-CoV-2 induced CPE for 4 days using an inverted optical microscope. The virus titer was estimated according to Spearman-Karber formula (Kundi, 1999) and defined as the reciprocal of the highest viral dilution leading to at least 50% CPE in inoculated wells.

### SARS-CoV-2 authentic virus neutralization assay

The neutralization activity of culture supernatants from monoclonal was evaluated using a CPE-based assay as previously described (Manenti et al., 2020). S-protein-specific memory B cells was initially evaluated by means of a qualitative live-virus based neutralization assay against a one-point dilution of the samples. Supernatants were mixed in a 1:3 ratio with a SARS-CoV-2 viral solution containing 25TCID_50_ of virus (final volume: 30 μl). After 1 hour incubation at 37°C, 5% CO_2_, 25 μl of each virus-supernatant mixture was added to the wells of a 96-well plate containing a sub-confluent Vero E6 cell monolayer. Following a 2-hour incubation at 37°C, the virus-serum mixture was removed and 100 μl of DMEM 2% FBS were added to each well. Plates were incubated for 3 days at 37°C in a humidified environment with 5% CO_2_, then examined for CPE by means of an inverted optical microscope. Absence or presence of CPE was defined by comparison of each well with the positive control (plasma sample showing high neutralizing activity of SARS-CoV-2 in infected Vero E6 cells) and negative control (human serum sample negative for SARS-CoV-2 in ELISA and neutralization assays). Following expression as full-length IgG1 recombinant antibodies were quantitatively tested for their neutralization potency against both the wild type and D614G strains. The assay was performed as previously described but using a viral titer of 100TCID50. Antibodies were prepared at a starting concentration of 20 μg/mL and diluted step 1:2. Technical triplicates were performed for each experiment.

### Characterization of SARS-CoV-2 RBD-Antibodies binding by Flow cytometry

Flow cytometry analysis was performed to define antibodies interaction with S-protein-receptor-binding domain (RBD). Briefly, APEX™ Antibody Labeling Kits (Invitrogen) was used to conjugate 20 μg of selected antibodies to Alexa fluor 647, according to the manufacturer instructions. Then, 1 mg of magnetic bead (Dynabeads™ His-Tag, Invitrogen) were coated with 70 μg of histidine tagged RBD. To assess the ability of each antibody to bind the RBD domain, 20 μg/mL of labelled antibody were incubated with 40 μg/mL of beads-bound RBD for 1 hour on ice. Then, samples were washed with 200 μL of Phosphate-buffered saline (PBS), resuspended in 150 μL of PBS and assessed with a FACSCanto II flow cytometer (Becton Dickinson). Results were analyzed by FlowJo (version 10).

### Affinity evaluation of SARS-CoV-2 neutralizing antibodies

Anti-Human IgG Polyclonal Antibody (Southern Biotech 2040-01) was immobilized via amine group on two flow cells of a CM5 sensor chip. For the immobilization, anti-human IgG Ab diluted in 10mM Na acetate pH 5.0 at the concentration of 25 μg/ml was injected for 360 sec over the dextran matrix, which had been previously activated with a mixture of 0.1M 1-ethyl-3(3-dimethylaminopropyl)-carbodiimide (EDC) and 0.4M N-hydroxyl succinimide (NHS) for 420 sec. After injection of the antibody, Ethanolamine 1M was injected to neutralize activated group. 10 μl/min flow rate was used during the whole procedure. Anti-SPIKE protein human mAbs were diluted in HBS-EP+ (Hepes 10 mM, NaCl 150 mM, EDTA 3.4 mM, 0.05% p20, pH 7.4) and injected for 120 sec at 10 μl/min flow rate over one of the two flow cells containing the immobilized Anti-Human IgG Antibody, while running buffer (HBS-EP+) was injected over the other flow cell to be taken as blank. Dilution of each mAb was adjusted in order to have comparable levels of RU for each capture mAb. Following the capture of each mAb by the immobilized anti-human IgG antibody, different concentrations of SPIKE protein (20 μg/ml, 10 μg/ml, 5 μg/ml, 2.5 μg/ml and 1 μg/ml in HBS-EP+) were injected over both the blank flow cell and the flow cell containing the captured mAb for 180 sec at a flow rate of 80 μl/min. Dissociation was followed for 800 sec, regeneration was achieved with a pulse (60 sec) of Glycine pH 1.5. Kinetic rates and affinity constant of SPIKE protein binding to each mAb were calculated applying a 1:1 binding as fitting model using the Bia T200 evaluation software 3.1.

## SUPPLEMENTARY FIGURES AND TABLES

**Table S1.** Identification of neutralizing antibodies. The Table shows numbers and percentages of neutralizing, partially neutralizing and not-neutralizing antibodies identified for each donor assessed in this study.

| Subject ID | S-protein<sup>+</sup> mAbs (ELISA) | Neutralizing mAbs (%) | Partially neutralizing mAbs (%) | Not-neutralizing mAbs (%) |
| --- | --- | --- | --- | --- |
| PT-004 | 158 | 6 (3,8) | 7 (4,4) | 145 (91,8) |
| PT-005 | 33 | 2 (6,1) | 2 (6,1) | 29 (87,9) |
| PT-006 | 34 | 3 (8,8) | 1 (2,9) | 30 (88,2) |
| PT-008 | 148 | 28 (18,9) | 20 (13,5) | 100 (67,2) |
| PT-009 | 61 | 9 (14,8) | 3 (4,9) | 49 (80,3) |
| PT-010 | 39 | 2 (5,1) | 1 (2,6) | 36 (92,3) |
| PT-012 | 132 | 8 (6,1) | 6 (4,5) | 118 (89,4) |
| PT-014 | 165 | 41 (24,8) | 12 (7,3) | 112 (67,9) |
| PT-041 | 165 | 28 (17,0) | 49 (29,7) | 88 (53,3) |
| PT-100 | 230 | 37 (16,1) | 22 (9,6) | 171 (74,3) |
| PT-101 | 219 | 49 (22,4) | 17 (7,8) | 153 (69,9) |
| PT-102 | 145 | 32 (22,1) | 10 (6,9) | 103 (71,0) |
| PT-103 | 129 | 34 (26,4) | 20 (15,5) | 75 (58,1) |
| PT-188 | 72 | 2 (2,8) | 2 (2,8) | 68 (94,4) |
| **Total (N)** | 1,731 | 281 | 172 | 1,278 |
| **Total (%)** |  | 16,2 | 10,0 | 73,8 |

**Table S2.** S-protein binding distribution of neutralizing antibodies. The Table summarizes the binding distribution of neutralizing antibodies against the S1- domain, S2-domain and S-protein trimer.

| Subject ID | SARS-CoV-2 Neutralizing mAbs | S1-domain specific (%) | S2-domain specific (%) | S-protein trimers specific (%) |
| --- | --- | --- | --- | --- |
| PT-004 | 13 | 10 (76,9) | 1 (7,7) | 2 (15,4) |
| PT-005 | 4 | 4 (100,0) | 0 (0,0) | 0 (0,0) |
| PT-006 | 4 | 3 (75,0) | 0 (0,0) | 1 (25,0) |
| PT-008 | 48 | 30 (62,5) | 8 (16,7) | 10 (20,8) |
| PT-009 | 12 | 2 (16,7) | 0 (0,0) | 10 (83,3) |
| PT-010 | 3 | 3 (100,0) | 0 (0,0) | 0 (0,0) |
| PT-012 | 14 | 9 (64,3) | 0 (0,0) | 5 (35,7) |
| PT-014 | 53 | 32 (60,4) | 6 (11,3) | 15 (28,3) |
| PT-041 | 77 | 47 (61,0) | 8 (10,4) | 22 (28,6) |
| PT-100 | 59 | 22 (37,3) | 5 (8,5) | 32 (54,2) |
| PT-101 | 66 | 31 (47,0) | 7 (10,6) | 28 (42,4) |
| PT-102 | 42 | 18 (42,9) | 6 (14,3) | 18 (42,9) |
| PT-103 | 54 | 33 (61,1) | 12 (22,2) | 9 (16,7) |
| PT-188 | 4 | 0 (0,0) | 0 (0,0) | 4 (100,0) |
| Total (N) | 453 | 244 | 53 | 156 |
|  | Total (%) | 57,5 | 7,3 | 35,2 |

**Table S3.** Potency distribution of SARS-CoV-2 S-protein specific nAbs. The Table reports the number of recombinant nAbs expressed per each subject and their distribution based on the neutralization potency.

| Subject ID | Expressed nAbs | Weakly Neutralizing mAbs (>500 ng/mL) | Medium Neutralizing mAbs (100- 500 ng/mL) | Highly Neutralizing mAbs (10- 100 ng/mL) | Potently Neutralizing mAbs (1 - 10 ng/mL) |
| --- | --- | --- | --- | --- | --- |
| PT-004 | 11 | 8 | 1 | 1 | 1 |
| PT-005 | 4 | 4 | 0 | 0 | 0 |
| PT-006 | 0 | 0 | 0 | 0 | 0 |
| PT-008 | 32 | 18 | 7 | 7 | 0 |
| PT-009 | 11 | 8 | 3 | 0 | 0 |
| PT-010 | 3 | 3 | 0 | 0 | 0 |
| PT-012 | 6 | 5 | 1 | 0 | 0 |
| PT-014 | 23 | 9 | 14 | 0 | 0 |
| PT-041 | 35 | 21 | 6 | 7 | 1 |
| PT-100 | 20 | 12 | 6 | 1 | 1 |
| PT-101 | 29 | 20 | 7 | 2 | 0 |
| PT-102 | 21 | 16 | 3 | 2 | 0 |
| PT-103 | 24 | 20 | 4 | 0 | 0 |
| PT-188 | 1 | 1 | 0 | 0 | 0 |
| Total (N) | 220 | 145 | 52 | 20 | 3 |
| Total (%) |  | 65,9 | 23,6 | 9,1 | 1,4 |

**Fig. S1.**
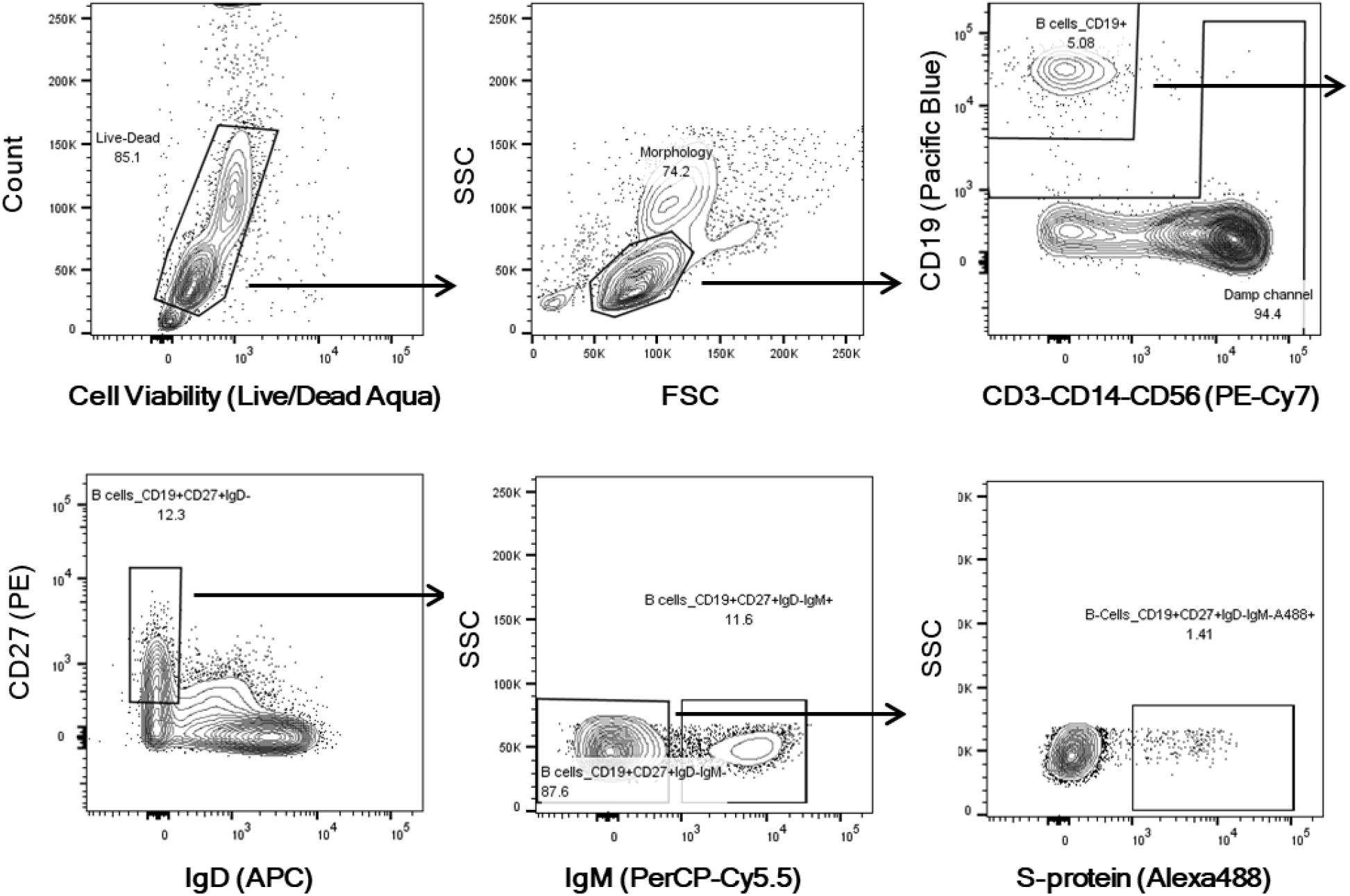
Gating strategy for S-protein specific MBC single cell sorting. Starting from top left to the right panel, the gating strategy shows: Live/Dead; Morphology; CD19+ B cells; CD19+CD27+IgD-; CD19+CD27+IgD-IgM-; CD19+CD27+IgD-IgM-S-protein+B cells.

**Fig. S2.**
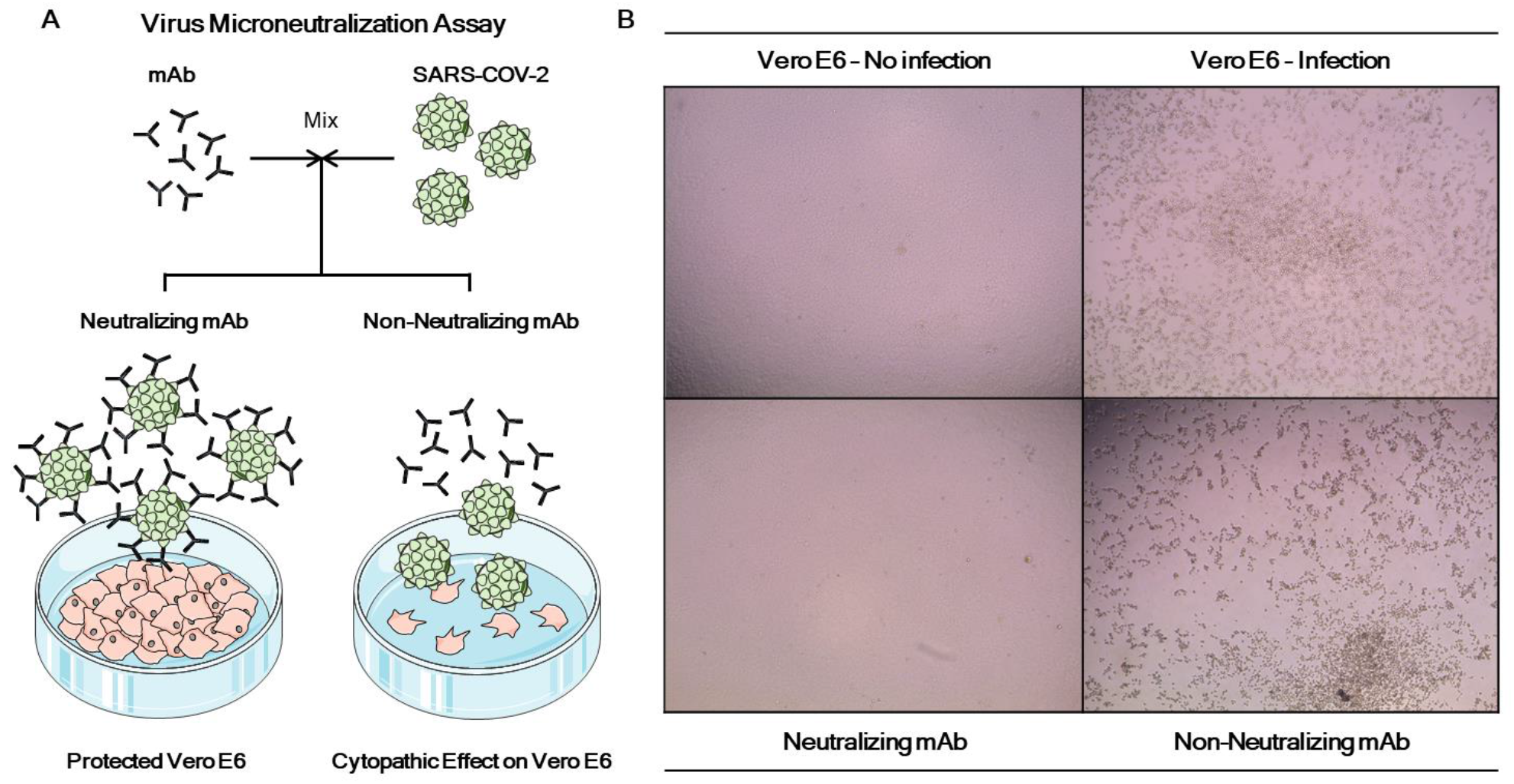
SARS-CoV-2 neutralization assay for S-protein specific mAbs. (A) Schematic representation of the virus neutralization assay used in this study to assess functional activities of S-protein specific mAbs. (B) Representative microscope images showing the cytopathic effect of SARS-CoV-2 or the protective efficacy of the screened supernatants on Vero E6 cells.

**Fig. S3.**
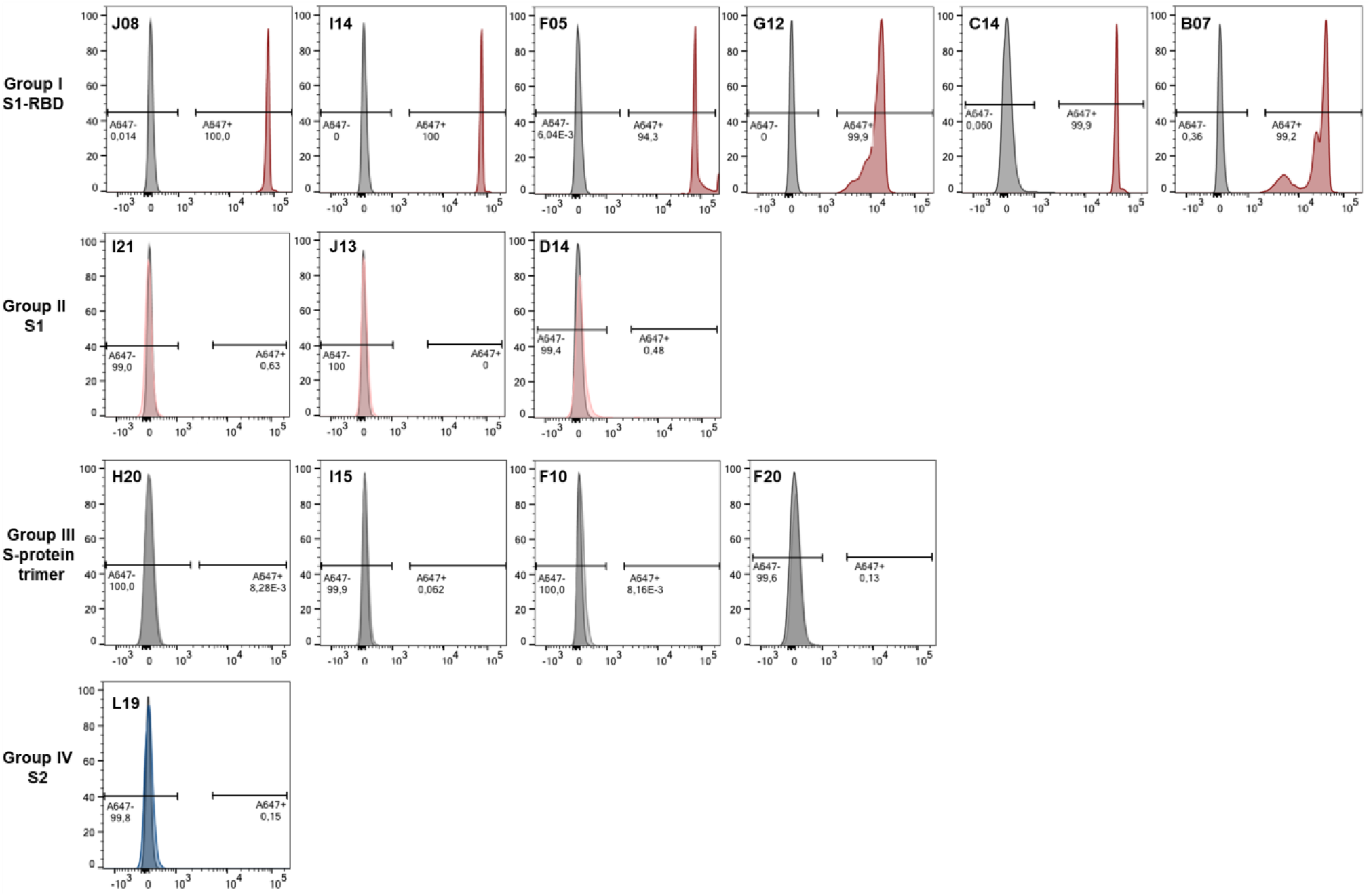
Binding to S-protein receptor binding domain. Histograms show the binding ability of selected antibodies to the S-protein RBD. Gray histogram represent the negative control while colored histograms show tested antibodies. Percentage of positive and negative populations are denoted on each graph.

**Fig S4.**
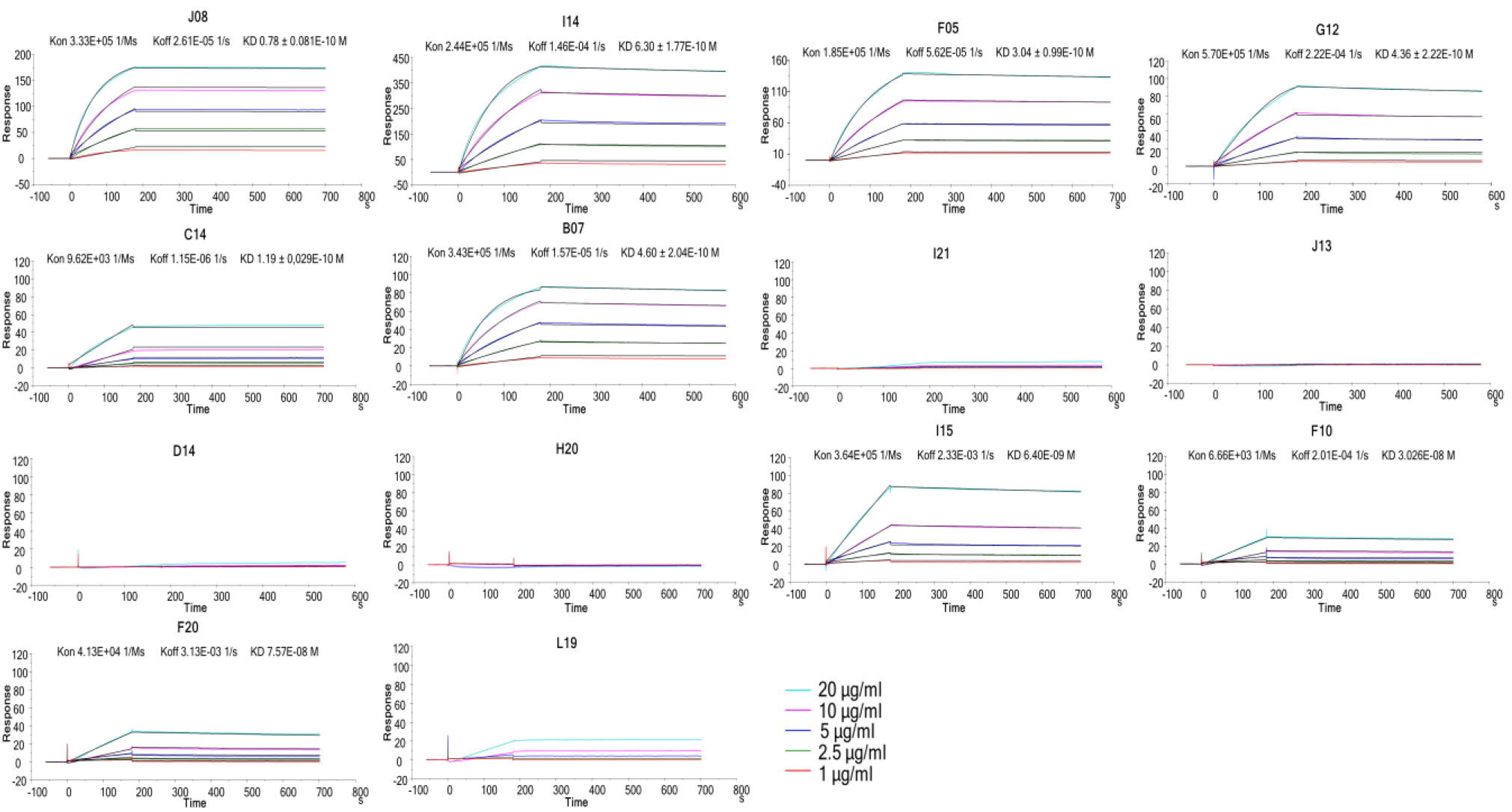
Binding kinetics of SARS-CoV-2 nAbs to the S-protein antigen. Representative binding curves of selected antibodies to SARS-CoV-2 S-protein trimer. Different curve colors define the spike concentration used in the experiment. Kon, Koff and KD are denoted on each graph.

